# Selective activation of alternative *MYC* core promoters by Wnt-responsive enhancers

**DOI:** 10.1101/266387

**Authors:** Jorge A. Bardales, Evin Wieser, Hideya Kawaji, Yasuhiro Murakawa, Xavier Darzacq

## Abstract

In Metazoans, transcription of most genes is driven through the use of multiple alternative promoters. Although the precise spatio-temporal regulation of alternative promoters is important for proper gene expression, the mechanism that mediates their differential utilization remains unclear. Here, we investigate how the two alternative promoters (P1, P2) that drive *MYC* expression are regulated. We find that P1 and P2 can be differentially regulated across cell-types, and that their selective usage is largely mediated by distal regulatory sequences. Moreover, we show that in the colon carcinoma cell line HCT-116, Wnt-responsive enhancers preferentially upregulate transcription from the P1 promoter using both transgenic reporter assays and in the context of the endogenous *Myc* locus upon Wnt induction. In addition, multiple enhancer deletions using CRISPR/Cas9 corroborate the regulatory specificity of P1. Finally, we show that preferential activation between Wnt-responsive enhancers and the P1 promoter is influenced by distinct core promoter elements present in the two *MYC* promoters. Taken together, our results provide new insights into how enhancers can specifically target alternative promoters and suggest that formation of these selective interactions could allow more diverse combinatorial regulation of transcription initiation.

## INTRODUCTION

RNA Polymerase II-dependent transcription initiation is a complex process that must be precisely regulated for the proper generation of spatio-temporal patterns of gene expression. The core promoter plays an essential function in integrating multiple transcriptional signals, including tissue-specific enhancer mediated activation to modulate the rate of transcription initiation. (Danino et al., 2015; Kadonaga, 2012; Ngoc et al., 2017). The core promoter can be defined as a short stretch of DNA surrounding the transcription start site (TSS) that potentiates directional transcription initiation. Core promoters are diverse in their composition of cis-control DNA and can contain multiple elements like TFIIB recognition sites (BRE^u^ & BRE^d^), TATA box binding sites, initiator sequences (Inr), motif ten elements (MTE) and downstream core promoter elements (DPE) (Lenhard et al., 2012; Ngoc et al., 2017). These diverse core promoter elements can serve two roles: they can modulate the basal activity of core promoters (Müller and Tora, 2014; Nogales et al., 2017), and they can regulate the core promoter capacity to be activated by specific enhancers and other distal cis-regulatory sequences (Butler and Kadonaga, 2001; Davis and Schultz, 2000; Juven-Gershon et al., 2008; Zabidi et al., 2015)

Numerous reports have shown that in metazoans, most genes are under the control of multiple alternative promoters (Carninci et al., 2006; Davuluri et al., 2008; Singer et al., 2008; Sun et al., 2011). In fact, the most recent report from the FANTOM consortium, which characterized more than 900 human cell samples by using Cap Analysis of Gene Expression (CAGE) found that genes, on average, can utilize 4 robust TSSs (The FANTOM Consortium and the RIKEN PMI and Clst (dgt), 2014). Interestingly, alternative promoters have been shown to be differentially utilized across different tissues (Carninci et al., 2006; Steinthorsdottir et al., 2004), during development and upon specific stimuli (Arner et al., 2015; Haberle et al., 2014), suggesting that fine tuning of alternative promoters may be important for proper gene expression. Importantly, mis-regulation of alternative core promoter usage can lead to pathological states including cancer and other disorders (Agarwal et al., 1996; Pedersen et al., 2002; Tan et al., 2007). Although initial evidence suggested that promoter usage is largely dependent on the epigenetic landscape of chromatin (Angeloni et al., 2007; Turner et al., 2008) and the presence of proximal transcription factor binding sites (Archey et al., 1999; Ngondo and Carbon, 2014; Pozner et al., 2007; Vitezic et al., 2010), the specific mechanisms that regulate alternative promoter usage remains poorly understood.

*MYC* is a key transcription factor that regulates the expression of thousands of genes involved in a wide-range of key cellular processes including cell growth, proliferation and metabolism (Conacci-Sorrell et al., 2014; Dang et al., 2006). Given the important physiological role of *MYC*, it is not surprising to find that MYC expression needs to be tightly regulated, especially at the transcription level (Pelengaris et al., 2002). Two alternative tandem promoters, P1 and P2, that are closely located drive *MYC* expression. These two promoters integrate multiple regulatory signals, including hundreds of tissue-specific enhancers (Fulco et al., 2016; Rennoll and Yochum, 2015; Uslu et al., 2014; Zhang et al., 2016), to precisely regulate *MYC* transcription. In this report, we find by analyzing CAGE data that the two *MYC* promoters can be differentially regulated across different human cell samples. Importantly, using artificial reporter assays, we found that the activity of the *MYC* promoters imbedded in transgenic constructs cannot recapitulate the differential promoter usage observed in the context of the endogenous Myc locus, suggesting that promoter usage is likely regulated by distal regulatory elements not present in the short synthetic constructs. This led us to test in colon carcinoma cells if distal *MYC* enhancers that are responsive to Wnt signaling could differentially regulate the endogenous activity of the two promoters. We observed that Wnt-responsive enhancers preferentially activate the P1 promoter upon Wnt induction. In addition, by using CRISPR/Cas9, we confirmed that enhancer deletion preferentially downregulates activity of the P1 promoter. Finally, we demonstrate that preferential activation of the P1 promoter is mediated by distinct promoter elements associated with P1 that differ from P2. Taken together, these results suggest that alternative promoters can mediate cell-type selective transcription regulation by facilitating specific promoter-enhancer interactions.

## RESULTS

### *MYC* alternative promoters can be differentially regulated

To gain a better understanding of how the two alternative *MYC* promoters might be differentially deployed, we surveyed P1 and P2 promoter usage across 869 human cell types and tissues from the FANTOM database (The FANTOM Consortium and the RIKEN PMI and Clst (dgt), 2014) **(FIGURE 1A)**. We counted the number of CAGE reads coming from each promoter within a 11bp window to calculate the P2 to P1 promoter usage ratio and its distribution across cell lines **(FIGURE 1B)**. We observed that the P2 promoter preferentially drives transcription of the *MYC* gene across different cell types in agreement with previous reports (Wierstra and Alves, 2008). Importantly, this analysis showed that the P2 to P1 promoter usage ratio distribution across the cell lines was very broad, spanning over two orders of magnitude (P2/P1_CAGE_= [0.37 to 34.24]). This suggested that transcription networks could differentially regulate the activity of the two tandem *MYC* promoters. Interestingly, inspection of the most extreme values showed that the lowest P2/P1_CAGE_ ratio observed was in the Burkitt’s lymphoma cell line RAJI. This cell line bears a translocation that has repositioned the switch region of the gamma heavy chain locus upstream from the *MYC* gene. This observation suggested that distal regulatory sequences may be important for differential core promoter usage.

**Figure 1.**
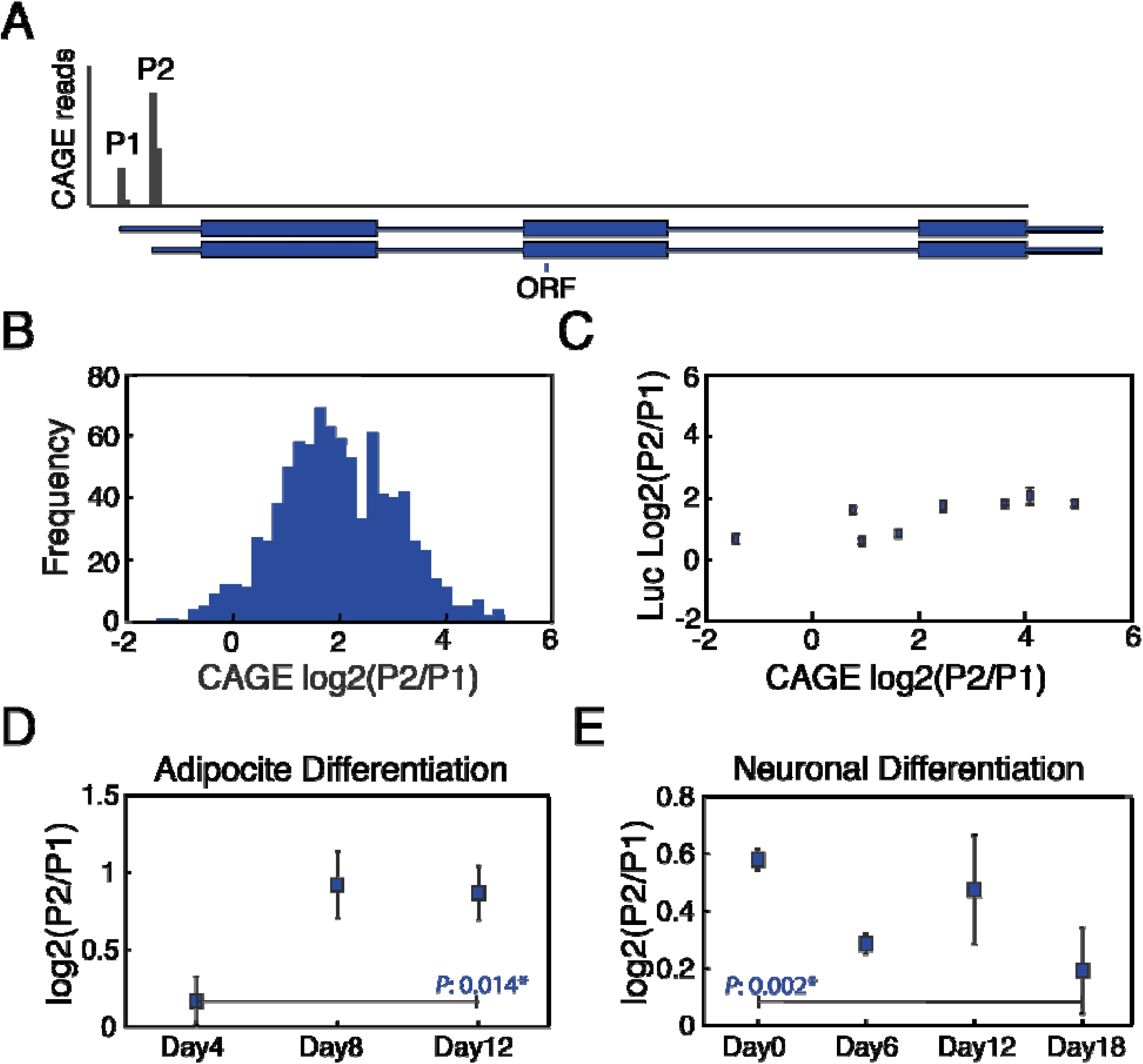
MYC transcription is driven by two alternative promoters which are differentially regulated in vivo.

(A) CAGE reads show that MYC gene is expressed by two alternative promoters, P1 and P2.
(B) Histogram of promoter usage ratio across 869 samples obtained from the FANTOM5 database.
(C) In vivo promoter usage ratio cannot be explained by promoter activity alone.
(D) Adipocyte differentiation leads to stronger usage of the P2 promoter.
(E) Neuronal differentiation leads to stronger usage of the P1 promoter

Based on these preliminary observations, we investigated if the in vivo promoter usage could be explained by differential promoter activation as a consequence of enhancer bias derived from the transcriptional network. For this we measured the independent activity of the P1 and P2 promoters by determining the activity of luciferase constructs harboring only proximal core P2 or P1 promoter sequences across eight cell lines with different P2/P1_CAGE_ ratios. This allowed us to calculate the synthetic P2 to P1 promoter activity ratio across multiple cell lines relative to endogenous promoter usage in the context of intact chromosomes **(FIGURE 1C)**. These results showed that in all cell lines tested, the P2 promoter displayed a higher basal activity (P2/P1_LUC_>1) and that this ratio is maintained across different cell lines (P2/P1_LUC_= [1.8,4.4]), especially when compared to the endogenous CAGE values. This result suggests that regulation of the 2 Myc promoters in transgenes does not recapitulate the pattern of differential P2/P1 usage observed for endogenous Myc in the context of native chromatin. This finding suggests that distal cis-regulatory elements are likely to be, at least in part, responsible for differential activation of Myc promoter P2 versus P1.

Next, we explored if changes in the transcriptional regulatory network during differentiation could lead to changes in alternative promoter usage. We analyzed 21 time courses of human cells exposed to differentiation cues and stimuli from the FANTOM database (Arner et al., 2015). In 7 out of these 21 time courses we observed a significant change in the P1/P2 promoter usage along the time course, suggesting that changes in the transcription regulatory network influenced differential promoter usage in a cell-type dependent manner. In addition, we observed that upregulation of the P1 versus P2 promoter usage was possible. For example, during Adipocyte differentiation, P2 promoter usage is stimulated **(FIGURE 1D)**, while during neuronal differentiation the P1 promoter activity is increased **(FIGURE 1E)**. This analysis suggests that transcription regulatory networks and distal enhancers could be responsible for differentially modulating the activation of the two alternate *MYC* promoters.

### Wnt-responsive enhancers preferentially activate the P1 promoter

To investigate if distal regulatory elements could specifically regulate the differential usage of the alternative P1 and P2 promoters, we tested how enhancers activate these 2 *MYC* promoters. We chose to work with the HCT-116 colon carcinoma cell line where multiple Wnt-responsive enhancers have been reported to regulate MYC expression (Rennoll and Yochum, 2015). These Wnt-responsive enhancers (shaded in red) are located throughout the MYC locus as can be seen based on their TCF7L2, RNA Polymerase II and H3K27Ac ChIP-Seq profiles **(FIGURE 2A)**. To test how the Wnt-responsive enhancers regulate the activity of the two alternative *MYC* promoters, we induced the Wnt pathway by treating the cells with different concentrations of LiCl (Shah et al., 2015) which promotes the relocalization of β-catenin and the subsequent activation of Wnt target genes **(FIGURE 2B)**. We observed that Wnt-activation increased the overall levels of *MYC* mRNA in a LiCl dependent manner **(Sup figure 1)**. More interestingly, we found that transcription from the P1 promoter is preferentially activated **(Figure 2C**), whereas transcription from P2 was not affected significantly **(Figure 2D).** This result suggests that Wnt induction preferentially activates transcription from the P1 promoter in HCT-116 cells.

**Figure 2.**
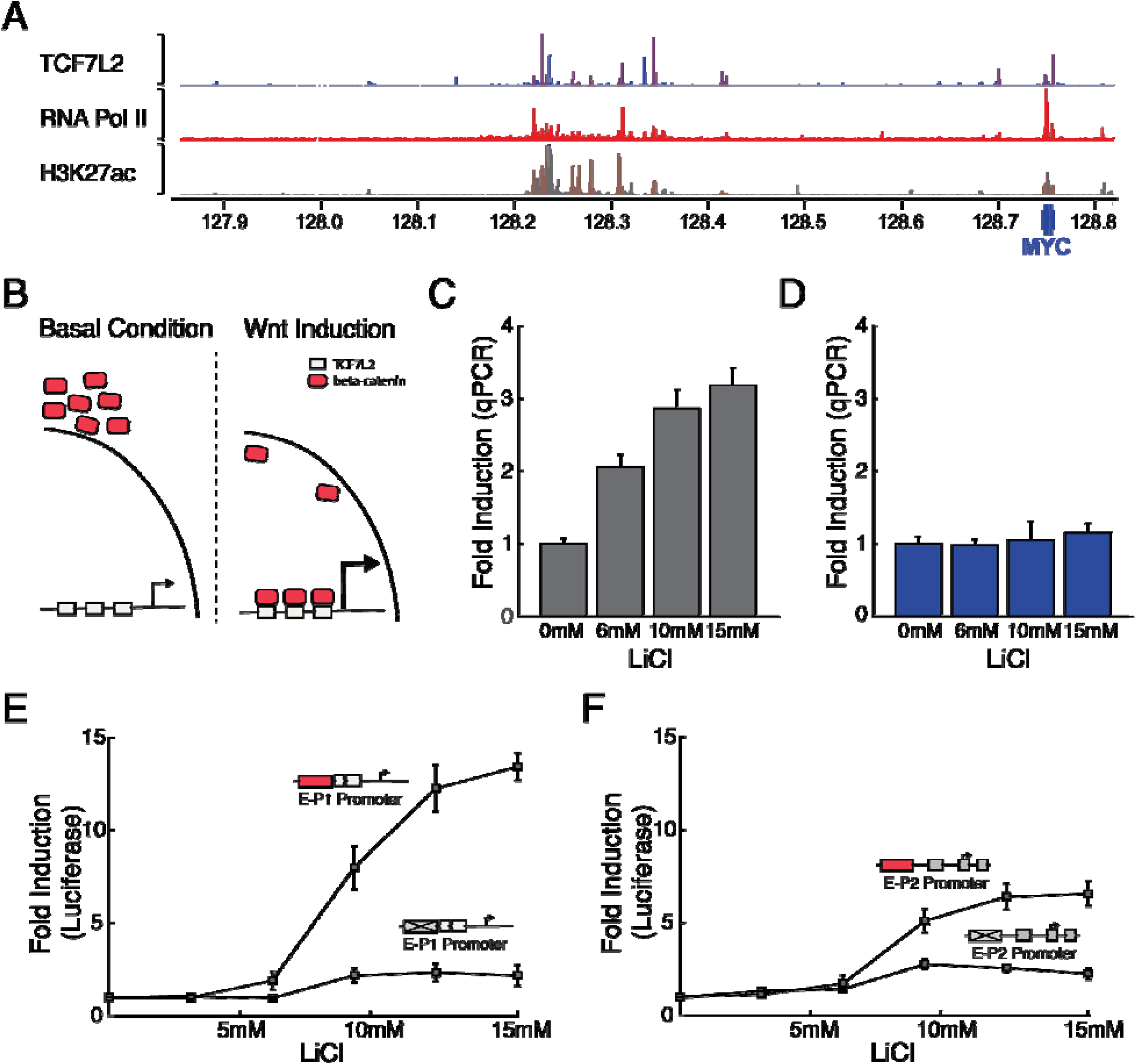
In HCT-116 cells, activation of Wnt-responsive enhancers preferentially upregulates the P1 promoter.

(A) Multiple Wnt-responsive enhancers (shaded in red) regulate MYC transcription. ChIP-seq traces of TCF7L2, RNA Pol II and H3K27ac mark the location of Wnt-responsive enhancers.
(B) Wnt Induction promotes relocalization of β-catenin.
(C) Wnt Induction strongly upregulates initiation from the P1 promoter.
(D) Wnt Induction does not upregulate initiation from the P2 promoter.
(E) Induction of Wnt-responsive enhancer strongly activates transcription of the P1 promoter
(F) Induction of Wnt-responsive enhancer midly activates transcription of the P1 promoter

To further probe how Wnt-responsive enhancers regulate the *MYC* promoters, we developed a system to independently quantify fold activation for each promoter by Wnt-responsive enhancers. We generated two luciferase reporter constructs with minimal P1 or P2 promoter under the regulation of a previously characterized strong Wnt-responsive enhancer (Veeman et al., 2003). We observed that the P1 promoter harboring enhancers was activated 12 fold when compared to the control P1 promoter with a mutated enhancer **(Figure 2E)**. By contrast, the P2 promoter was activated 6 fold in comparison to its own control **(Figure 2F)**.

### Enhancer deletions preferentially downregulate transcription from the P1 promoter

Next, we investigated the effect of deleting Wnt-responsive enhancers on the two alternative *MYC* promoters. Five enhancers were selected across the topological domain located at approximately 1 Kb, 7 Kb, 335 Kb, 405 Kb and 550 Kb distal to the *MYC* gene body. This set of enhancers had been previously reported to show strong interactions with the gene body as determined by Hi-C capture in colon carcinoma cells (Jäger et al., 2015), and were named respectively E1 to E5. We next used CRISPR/Cas9 to generate cell lines with homozygous enhancer deletions **(Figure 3A)**. Here, sequence-specific guide RNAs were designed flanking each enhancer to generate homozygous enhancer deletions surrounding the TCF7L2 binding sites **(Sup Figure 2)**. When we tested the effect of these enhancer deletions on *MYC* mRNA, we found that in four cases the deletions caused significant downregulation of total *MYC* mRNA levels **(Figure 3B)**. Downregulation of *MYC* levels was accompanied by defects in cell growth **(Sup Figure 3),** suggesting that these enhancers are important for *MYC* regulation across the cell cycle. When the specific activity of the two alternative *MYC* promoters was measured, we observed that the enhancer deletions preferentially downregulated transcription initiation from the P1 promoter **(Figure 3C),** while we only observed a minor reduction of transcription initiation from the P2 promoter **(Figure 3D)**. In sum, these genome editing experiments further support a preferential regulatory relationship between a select set of Wnt-responsive enhancers and the P1 promoter of Myc.

**Figure 3.**
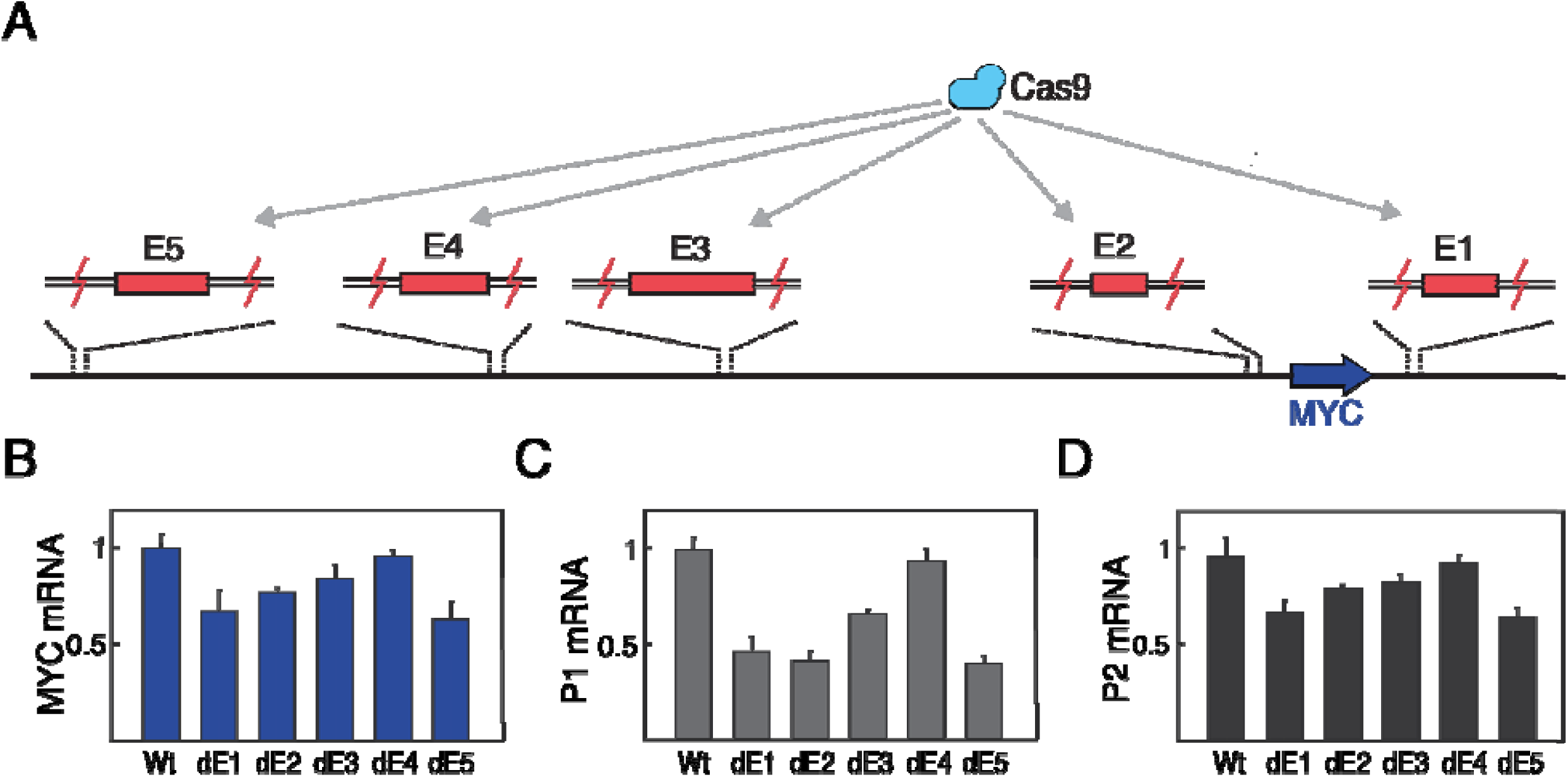
Wnt-responsive enhancer deletions preferentially downregulate transcription from the P1 promoter.

(A) Five enhancers were selected across the MYC locus to be deleted by using CRISPR/Cas9.
(B) Effect of enhancer deletion on MYC mRNA levels
(C) Enhancer deletions strongly downregulate transcription from the P1 promoter.
(D) While in the P2 promoter the enhancer deletions only cause minor downregulation

### Distinct core promoter architecture influences specific enhancer-promoter communication

Previous studies have shown that enhancer-promoter specificity can be mediated by the presence or absence of different core promoter elements (Juven-Gershon et al., 2008; Zabidi et al., 2015). To determine whether core promoter architecture comprised of specific elements might also mediate the selectivity of Wnt-responsive enhancers, we analyzed the promoter architecture of the two *MYC* core promoters. We found that the two alternative *MYC* core promoter possess rather distinct composition and disposition of their promoter elements **(Figure 4A).** Importantly, the type and disposition of promoter elements present in the two distinct *MYC* promoters are highly conserved in mammals **(Sup. Figure 4)**, suggesting the arrangement of elements in these 2 dissimilar promoter architectures could be important for function.

**Figure 4.**
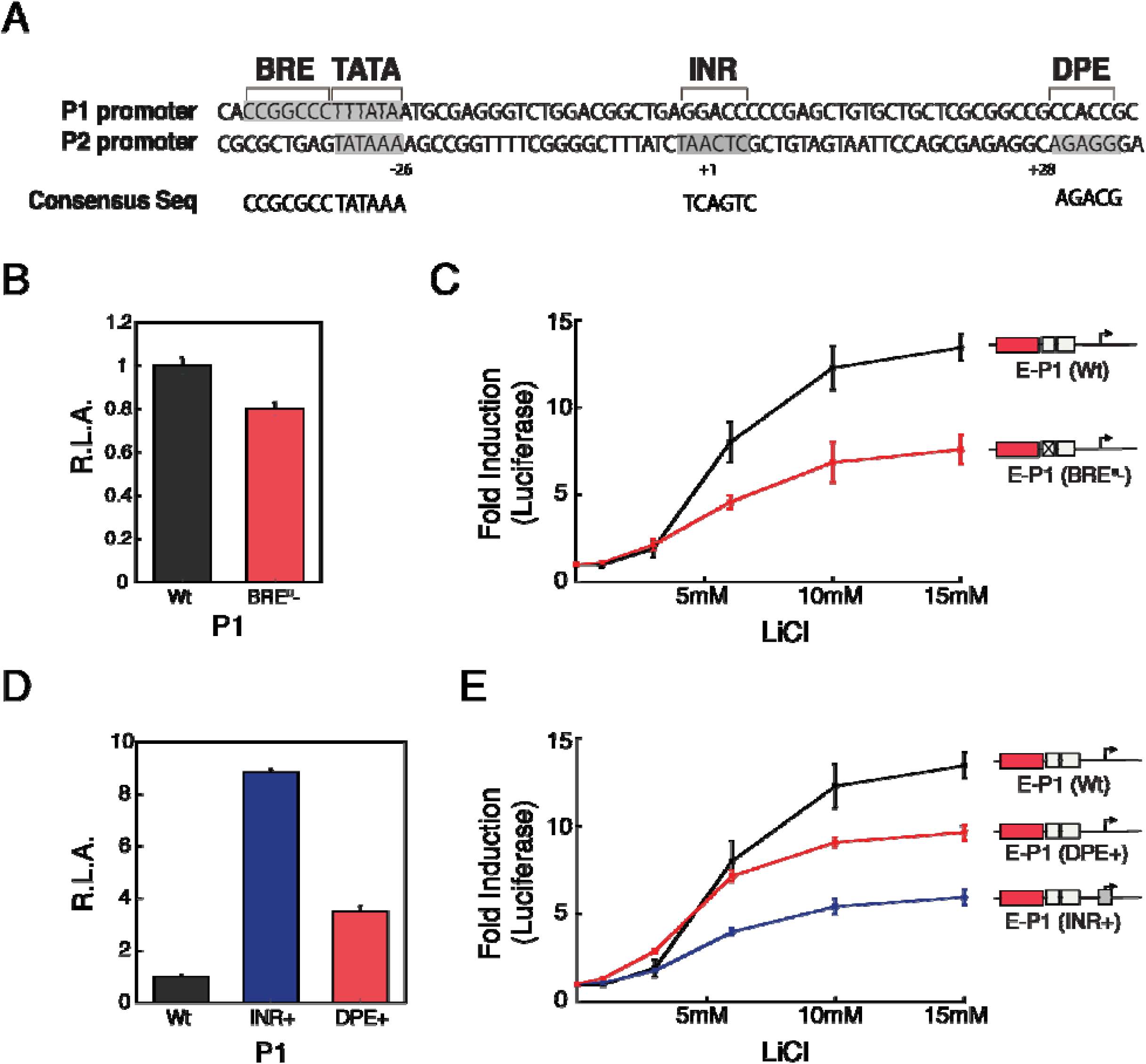
Promoter architecture mediates differential enhancer communication.

(A) MYC Promoter possess dissimilar promoter architecture. While the P1 promoter has a TATA box and a BRE^UP^ motif, the P2 promoter possess a strong TATA box, a INR and a DPE motif.
(B) P1 promoter lacking the BRE^u^ motif decreased minimally the basal activity.
(C) P1 promoter lacking the BRE^u^ motif had a drastic effect on the maximum fold activation.
(D) P1 promoter with either a consensus INR or DPE motif have stronger basal activity.
(E) The addition of a consensus INR or DPE motif to the P1 promoter diminishes fold activation.

In order to determine the basis for the preferential activation of P1 upon Wnt-induction, we investigated the role of the different promoter elements present within the two *MYC* core promoters. We first tested the role of the BRE^u^ on Wnt-dependent activation which is only present within the P1 promoter. We observed that mutation of the BRE^u^ motif in P1 had little effect on its basal promoter activity **(Figure 4B)**. In contrast, the presence of the BRE^u^ motif on the P1 promoter was crucial for Wnt-responsive enhancer-mediated activation **(Figure 4C)**. Furthermore, we observed that insertion of a consensus BRE^u^ motif into P2 had little effect on its basal activity, but increased its maximum fold activation when compared to the wild-type P2 promoter **(Sup. Figure 5).** These loss and gain of function experiments suggest that the BRE^u^ motif likely plays a role in potentiating the capacity of P1 to be differentially activated by Wnt-responsive enhancers.

Finally, we tested how 2 other core promoter elements, the INR or the DPE motifs, might influence the capacity of the 2 alternative promoters to be activated by the Wnt-responsive enhancers. First, we generated altered P1 promoter constructs containing consensus INR or DPE elements inserted into P1 which normally lacks such consensus sites. We observed that these altered P1 promoters carrying consensus INR or DPE motifs displayed stronger basal promoter activity compared to wild-type P1 **(FIGURE 4D)**. Interestingly, the presence of these 2 consensus motifs decreased the capacity of the P1 promoter to be activated by Wnt-responsive enhancers (**FIGURE 4E),** which is inversely correlated with the observed increase in its basal promoter activity. Conversely, disrupting strong INR or DPE elements that are normally present within the native P2 promoter resulted in an increased fold activation induced by the Wnt-enhancers accompanied by a proportional decrease in its basal promoter activity **(Sup. Figure 5).** These results suggest that the INR and DPE might dampen the fold activation of enhancer-induced transcription by elevating the basal promoter activity.

## DISCUSSION

The usage of alternative promoters as drivers of transcription initiation is an prevalent phenomenon that occurs throughout metazoans (Davuluri et al., 2008; Klerk and Hoen, 2015). Although numerous reports have shown that differential regulation of alternative promoter is important for proper development and cellular maintenance (Davuluri et al., 2008), the mechanisms that mediate alternative promoter usage remained largely unknown. Here, we show that selective enhancers can differentially regulate the activity of 2 core promoters driving the transcription of *MYC*. Specifically, we found that the *MYC* promoters, P1 and P2, are differentially deployed across different tissues and cell types and that distal regulatory elements are necessary for the targeted regulation of alternative core promoters. In addition, in HCT-116 colon carcinoma cells, we demonstrate that Wnt-responsive enhancers preferentially activate the P1 promoter. Finally, we show that the preferential activation of the P1 promoter is dependent on specific core elements and its promoter architecture revealing and important aspect of the mechanism directing alternative core promoter specificity.

Previous studies have reported that certain promoter elements present within the core promoter architecture are sufficient to specify enhancer-promoter communication (Butler and Kadonaga, 2001; Juven-Gershon et al., 2008; Zabidi et al., 2015). Here, we have extended these earlier studies by establishing that there is a preferential communication of select Wnt-responsive enhancers with certain elements present in the P1 promoter but absent in P2. Moreover, this selective promoter/enhancer communication is mediated by the combined action of multiple core promoter elements via two mechanisms. First, the presence of a BRE^u^ motif potentiates activation of P1 independent of its basal activity. And second, the presence of INR or DPE motifs apparently limits the range of enhancer dependent promoter activation by increasing the basal activity of the core promoter. This means that, in HCT-116 cells, the P2 promoter serves as a basal promoter largely insensitive to Wnt-responsive enhancers, whereas the P1 promoter can be fine-tuned by Wnt-responsive enhancers. The presence of these two differentially regulated promoters could be important for precise regulation of transcription initiation, especially of tightly regulated genes like *MYC*.

Multiple mechanisms are thought to regulate the specific formation of enhancer-promoter interactions that include chromosome topology, changes in the epigenetic chromatin landscape and biochemical compatibility (Arensbergen et al., 2014). The observation that small changes in alternative core promoter elements can differentially communicate with distal enhancers could further diversify the combinatorial specificity of these highly regulated interactions. Interestingly, genome-wide analysis of mammalian promoters have shown that the presence of alternative promoters is overrepresented in highly regulated genes (Baek et al., 2007). The MYC gene is exquisitely regulated at the transcriptional level, with more than 270 enhancers annotated in the enhancer atlas to regulate MYC, of which dozens have been functionally validated (Fulco et al., 2016; Sotelo et al., 2010; Uslu et al., 2014; Zhang et al., 2016). The differential usage of two alternative Myc core promoters would significantly expand the network of specific long distance enhancer/promoter interactions necessary to mediate different temporal and spatial transcriptional outcomes during development and differentiation.

## MATERIALS AND METHODS

### CAGE DATA ANALYSIS

*MYC* locus CAGE data was obtained from the FANTOM5 database (http://fantom.gsc.riken.jp/). The total number of counts was integrated within a 11bp window surrounding the TSS for each promoter and used to calculate the promoter ratio usage across 869 cell samples.

### CELL CULTURE

All cell lines used in this study were cultured in accordance with ATCC and Riken Cell bank guidelines in their suggested media supplemented with 10% fetal bovine serum in a humidified incubator at 5% CO2. For HCT-116 cells, Wnt induction was achieved by treating the cells with LiCl solution at different concentrations for 6hrs (Real time qPCRs) or 24hrs (luciferase reporter assays).

### REPORTER LUCIFERASE ASSAY

Luciferase reporter constructs were generated using pGL3-Basic luciferase reporter vectors. The *MYC* promoter sequences were cloned between XhoI and HindIII sites and the Wnt-responsive element was cloned between KpnI and NheI sites. Mutations of promoter elements was done using QuickChange II Protocol. The Primers used for site direct mutagenesis are listed in Table S1. The final mutant promoter sequences are in Annex S1. Reporter constructs were transiently transfected into cells using Lipofectamine 2000 along with a control plasmid (Renilla luciferase SV40) for 24 hours before luciferase activity was measured

### REAL TIME qPCRs

Total RNA was isolated using RNeasy Kit (QIAGEN), cDNA was generated using Maxima First Strand cDNA Synthesis Kit (Thermo Fisher) and qPCRs were done using SYBR FAST qPCR Mix (KAPA). In all cases manufacturer guidelines were followed. Primers used for quantitative RT-PCR were designed with Primer3 (http://bioinfo.ut.ee/primer3-0.4.0/) and are listed in Table S2.

### GENOME EDITING

To generate enhancer deleted HCT-116 clones, gRNAs were designed using the CRISPR design tool (http://crispr.mit.edu/) and cloned into a modified px330 vector (Cong et al., 2013). Two px330 vectors coding for gRNAs surrounding the enhancer locus were transfected into HCT-116 cells and posteriorly plated into 96 well plates. Generation of px330 plasmid was done using primers listed in in Tables S3.

## ACKNOWLEGEMENTS

We thank Gina Dailey for help with designing constructs and the Tjian-Darzacq Lab as well as the Doudna Lab for critical comments and discussion while preforming this Work. We thank Jennifer Doudna and Yoshihide Hayashizaki for their comments and feedback. We thank Robert Tjian for discussions and critical reading of the manuscript. We also want to thank Dr. Kartoosh Heydari at the Li Ka Shing Facility for flow cytometry assistance. This work was supported by NIH grant U54-DK107980 and by the California Institute of Regenerative Medicine grant LA1-08013 to XD.

## SUPPLEMENTAL INFORMATION

Supplemental Information includes supplemental Experimental Procedures, five figures, and four tables can be found with this article online.

